# Live-cell single-molecule dynamics of eukaryotic RNA polymerase machineries

**DOI:** 10.1101/2024.07.08.602478

**Authors:** Yick Hin Ling, Chloe Liang, Sixiang Wang, Carl Wu

## Abstract

Eukaryotic gene expression in the nucleus is orchestrated by three RNA polymerases (RNAP-I, -II, and -III) and associated factors^1,2^. Despite extensive biochemical, genomic, structural, and imaging studies, the real-time dynamics of these transcription complexes remain obscure. Here, we employ single-molecule tracking in living yeast to assess the physiological kinetics of over 50 representative proteins encompassing all three RNAP machineries. Components of RNAPI and RNAPIII pre-initiation complexes (PICs) engage in long-lived interactions on chromatin, reflecting their roles for constitutive rRNA and tRNA synthesis, in contrast to the transient RNAPII PIC. We further report the dynamics of key components across the RNAPII transcription cycle^2–5^—factors for upstream regulation, elongation, histone modification, RNAPII C-terminal domain (CTD) modification, RNA processing, and termination—revealing unprecedented insights into the temporal landscape of RNAPII transcription. Strikingly, many elongation factors, previously thought to travel processively with RNAPII, display transient residence times, suggesting highly dynamic interactions rather than constant association. Systematic screening of RNAPII-associated factors shows that truncation of RNAPII-CTD substantially reduces U1 snRNP residence time and decreases intron retention in ribosomal protein genes, providing insights into how CTD length influences co-transcriptional splicing. Our findings establish a framework for dynamic chromatin interactions of RNA polymerase machineries in living cells.

## Introduction

Eukaryotic transcription is accomplished by three nuclear RNA polymerases (RNAPs): RNAP-I, -II, and -III^1^. RNAPI synthesizes ribosomal RNAs (rRNAs), which are essential components of the ribosome. RNAPII transcribes protein-coding genes into messenger RNAs (mRNAs) and non-coding RNAs that regulate cellular processes^6,7^. RNAPIII synthesizes small RNAs, including 5S rRNA, an essential ribosome component, and transfer RNAs (tRNAs) for mRNA translation. Each polymerase complex operates with distinct factors modulating its activity at various stages of transcription, from initiation through elongation to termination^2^. Despite decades of extensive biochemical, genomic and structural investigations and recent super-resolution imaging studies, the temporal regulation of the three RNAP machineries and their regulatory factors on chromatin in living cells is far from complete. Important models, such as the formation of partial transcription pre-initiation complexes (PIC)^8,9^, RNAP recycling for repeated rounds of transcription^10,11^ and gene promoter-terminator looping^12^, are still being explored. Moreover, outstanding questions remain regarding which elongation factors interact transiently or bind persistently to the processively elongating RNAP^13^, and whether termination factors are recruited in an ad hoc manner or act continuously as pre-engaged transcription roadblocks^14^. In this study, we employ single-molecule tracking (SMT) in living yeast to systematically quantify the global physiological residence times of over 50 representative transcription proteins or protein complexes, covering key components of the multi-step transcription cycle for the three RNAPs in the nucleus, and establish a fundamental framework for the temporal dynamics of eukaryotic transcription. Our findings reveal distinct dynamics for each of the three RNAP PICs relevant to their specific functions, and further highlight a spectrum of long-lived and transient interactions for transcription elongation and termination factors.

The intrinsically disordered C-terminal domain (CTD) of RPB1 (Rpo21 in yeast), unique to the largest subunit of RNAPII, features 26 heptad repeats (YSPTSPS) in yeast and 52 repeats in mammals. The CTD is functionally modified mainly by phosphorylation^3–5^, resulting in a diverse array of binding sites for multiple CTD-interacting factors for regulation of transcription and co-transcriptional activities^15–18^. CTD truncation of RPB1 leads to transcription defects^18–29^ and reduces the size of liquid droplets or condensates formed by purified CTD *in vitro*^30,31^. In yeast, the CTD maintains diffusion confinement of RNAPII around active genes, facilitating its search for chromatin targets^32^. Here, we have compared the binding dynamics of CTD-associating factors in a viable but growth-impaired truncation mutant with 9 heptads (CTD9)^32^. We found minimal changes of residence times for most associating factors, but a sizeable reduction in the residence time of the U1 snRNP, with a functional decrease in intron retention for ribosomal protein genes, suggesting that CTD length can influence co-transcriptional splicing.

### Single-molecule tracking and refined modeling reveal residence times of RNAP-I, -II, and -III

We used SMT to measure the dynamics of representative subunits of RNAPI (Rpa190), RNAPII (Rpo21), and RNAPIII (Ret1) fused to HaloTag (Fig. 1a-c). A fast tracking imaging regime at high laser power and a short frame rate (10 ms/frame) captures both freely diffusing and chromatin-bound molecules (Fig. 1a-c and Supplementary Video 1). We classify particle trajectories as free or bound (Fig. 1c) based on diffusive characteristics, including the apparent diffusion coefficient (D) and the anomalous diffusion exponent^32^. The three nuclear RNA polymerases show bound trajectories at their expected subnuclear regions, visually demonstrating their spatially specific functions^32^ (Fig. 1c and Extended Data Fig. 3f). As illustrated in Figure 1c and corroborated by our previous study^32^, free RNAPII is predominantly concentrated in the nucleoplasm, with marked exclusion from the nucleolus. Similarly, free RNAPIII is also excluded from the nucleolus, albeit to a lesser extent. Interestingly, a detectable amount of free RNAPI is observed in the nucleoplasm, but its bound trajectories are mainly located in the nucleolus.

**Fig. 1.**
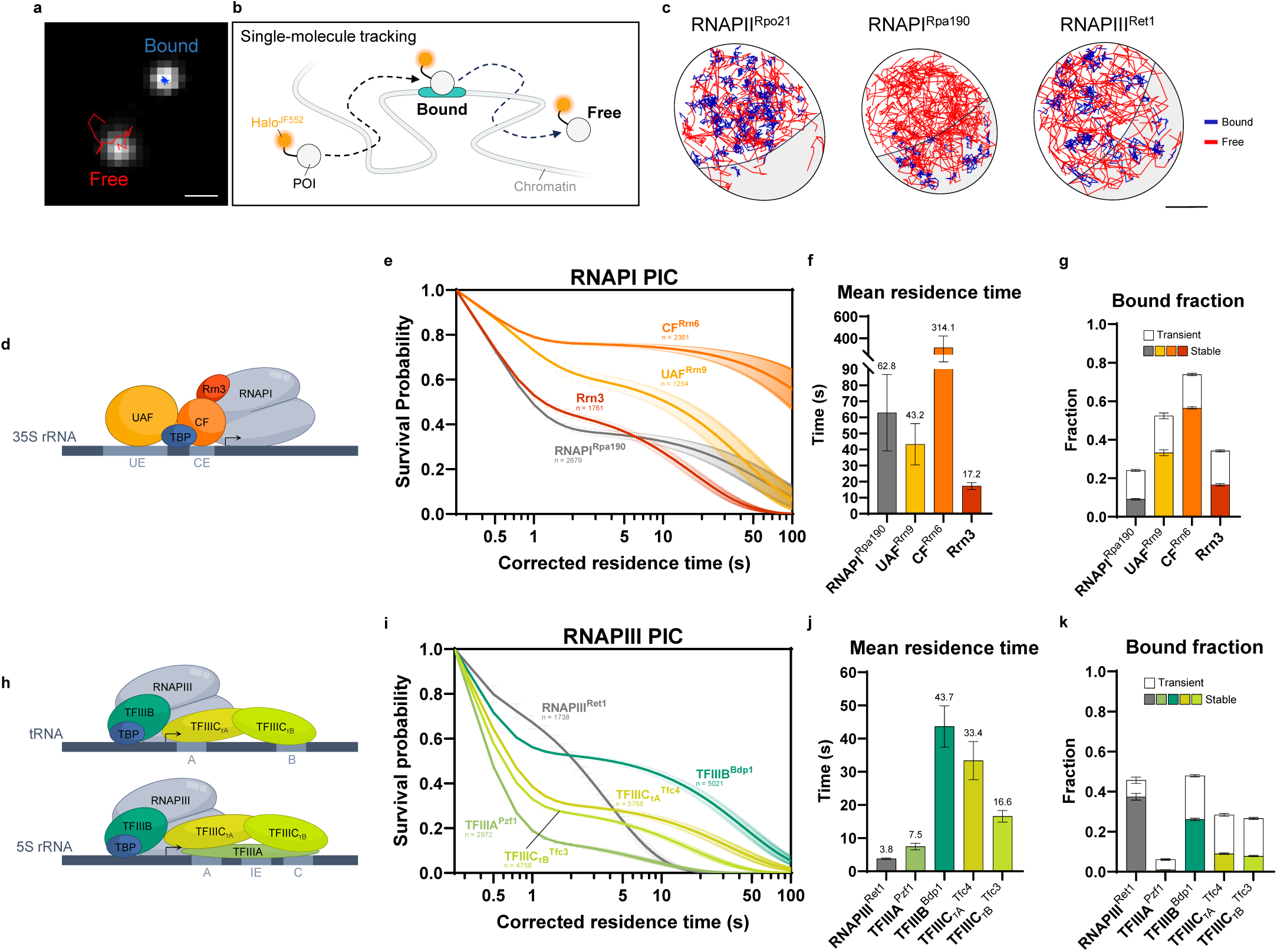
RNAPI and RNAPIII PICs exhibit prolonged stability. **a**, Example of bound and free single-molecule trajectories. Scale bar: 500 nm. **b**, Protein of interest (POI) fused with HaloTag and labelled with JF552 for single-molecule tracking. **c**, Bound and free trajectories of RNAP-II, -I, and -III in the nucleus captured by fast tracking. 100 random trajectories were shown respectively. Nucleolus highlighted in grey. Scale bar: 500 nm. **d**, Schematic of RNAPI PIC. UE = upstream element; CE = core element. Transcription start site (TSS) indicated by arrow. **e**, Survival probability of H2B-corrected residence times for RNAPI PIC (n = number of trajectories; mean value ± s.e.m.). **f**, Mean residence time of the stably bound fraction for RNAPI PIC (n = 5,000 resamplings; mean value ± s.e.m.). CF^Rrn6^ exhibited long residence time close to the H2B control limit; resamplings were thus constrained to yield non-negative decay constant for an approximated residence time. **g**, Transiently and stably bound fraction of RNAPI PIC (n = 5,000 resamplings; mean value ± s.e.m.). **h**, Schematic of RNAPIII PIC. A = A box; B = B box; IE = intermediate element; C = C box. TSS indicated by arrow. **i**, Survival probability of H2B-corrected residence times for RNAPIII PIC (n = number of trajectories; mean value ± s.e.m.). **j**, Mean residence time of the stably bound fraction for RNAPIII PIC (n = 5,000 resamplings; mean value ± s.e.m.). **k**, Transiently and stably bound fraction of RNAPIII PIC (n = 5,000 resamplings; mean value ± s.e.m.).

In addition to fast tracking, we utilized a slow tracking’ regime that employs low laser power to minimize photobleaching, and a long frame rate (250 ms/frame) to allow motion blur. This approach captures the binding kinetics of bound molecules, with minimal interference from the freely diffusing population (Extended Data Fig. 1c and Supplementary Video 2). We fit the survival probability of chromatin-bound molecules to a two-exponential decay model to dissect the fraction of transient and stable chromatin binding (Extended Data Fig. 2d-g and k-l), and estimated the global mean residence time of the stably bound molecules (hereafter residence time’; Extended Data Fig. 2d-j). Because the temporal resolution of slow tracking is limited by photobleaching, nuclear and chromatin movement, microscopic drift, and dye photophysics, we introduced a novel modeling approach by correcting the two-exponential decay curve against the stretched exponential decay recorded for long-lived histone H2B (2-exp_StrCORR_; Extended Data Fig. 2a-d and Methods). This refined correction improves the precision of the two-exponential fit for RNAP-I, -II, and -III (Extended Data Fig. 2e-i), revealing mean residence times of ∼60 s for RNAPI^Rpa190^, ∼40 s for RNAPII^Rpo21^ and ∼4 s for RNAPIII^Ret1^ (Fig. 1f,j, 2c and Extended Data Fig. 2e-j). These residence times, which reflect the global average duration of RNAPs binding on chromatin during initiation, elongation, and termination, correlates with the geometric mean of their respective gene lengths (Extended Data Fig. 1d-f), suggesting correspondence with functional activity.

### RNAP-I and -III PICs are long-lived, while RNAPII PIC is transient

RNAPI initiation in yeast begins with the binding of the 6-subunit upstream activation factor (UAF) to the upstream element (UE) of 35S rRNA gene^33^. UAF mediates the recruitment of the TATA-binding protein (TBP), a component common to all three nuclear RNAP PICs (Fig. 1d,h and 2a), which acts as a bridge to position the 3-subunit core factor (CF) to the core element (CE), located downstream of UE^33^. The binding of CF to CE allows subsequent recruitment of the Rrn3-bound RNAPI to form the complete RNAPI PIC (Fig. 1d), where Rrn3 facilitates the polymerase binding to the PIC components^2^. Upon initiation, RNAPI dissociates from Rrn3, CF, and UAF^34^, and the rest of the PIC potentially remains at the promoter^35–37^. We find a residence time of ∼40 s for UAF^Rrn9^, ∼20 s for Rrn3, and on the order of minutes for CF^Rrn6^ (Fig. 1e-g). These residence times of RNAPI PIC components are far longer than those observed in previous SMT studies of yeast RNAPII PIC components^38^ and other chromatin-binding factors, including chromatin remodelers^39^ and transcription factors^32,40,41^, which typically last for only several seconds. In yeast, the rRNA gene can be transcribed simultaneously by ∼50 RNAPI molecules^42^. Mechanistically, the observed long binding of the RNAPI PIC, especially for UAF and CF, supports a model in which these factors stay bound on the promoter, allowing efficient and persistent promoter priming for recruitment of multiple RNAPI complexes to yield a continuous chain of elongating polymerases^43^. *In vitro* studies suggest that only CF and Rrn3 are required for RNAPI basal transcription^44^, highlighting the importance of the long-lived CF for reinitiation of 35S rRNA synthesis.

RNAPIII transcription initiation at tRNA gene promoters (RNAPIII-type II promoters) requires TFIIIB and TFIIIC (Fig. 1h; top), with a third factor TFIIIA additionally required for 5S rRNA gene promoters (RNAPIII-type I promoters)^45^ (Fig. 1h; bottom). At tRNA promoters, the two DNA-binding subcomplexes of TFIIIC, *τ*A and *τ*B, bind respectively to the A and B box sequences, downstream of the transcription start site (TSS). At 5S rRNA gene promoters lacking the B box, recruitment of TFIIIC requires prior binding of TFIIIA to the A box, intermediate element (IE), and C box (Fig. 1h; bottom). TFIIIC binding assists in the recruitment and positioning of TFIIIB (TBP, Bdp1, and Brf1) to a region upstream of the TSS. Although TFIIIB can bind independently to the TATA element present upstream for some RNAPIII genes *in vitro*^46^, its engagement with chromatin in *vivo* requires TFIIIC^47^ (Fig. 1h). Structural studies have suggested that TFIIIB recruitment leads to the dissociation of TFIIIC, with the *τ*B subcomplex being displaced^48^. *τ*A may remain in contact with TFIIIB, followed by the recruitment of RNAPIII for transcription. Once TFIIIB binds stably on the promoter, it supports multiple rounds of RNAPIII transcription in *vitro*^49^.

Our live-cell SMT data extend the biochemical and structural perspective of RNAPIII transcription initiation, providing additional insights into the physiological timing of these processes (Fig. 1i-k). Among all RNAPIII PIC components, TFIIIB^Bdp1^ has the longest residence time of ∼45 s (Fig. 1j), consolidating its key role in directing RNAPIII reinitiation^49^. In line with the distinct dissociation of TFIIIC^48^, the residence time of TFIIIC*_τ_*_B_^Tfc3^ is half as long as TFIIIC*_τ_*_A_^Tfc4^ (16.6 s versus 33.4 s) (Fig. 1j). In contrast, TFIIIA^Pzf1^ has a residence time of 7.5 s (Fig. 1j), suggesting a more transient role in the assembly of the PIC for 5S rRNA genes. In addition to stable TFIIIB binding, recycling of the same RNAPIII molecule for multiple rounds of transcription has been observed *in vitro* to enhance transcription efficiency^50^. Assuming elongation dominates the observed 3.8 s residence time of RNAPIII^Ret1^ (Fig. 1j and Extended Data Fig. 1f), we estimate an elongation speed of 1.45 kb/min. This speed, comparable to previous *in vitro* estimates of 1.74 kb/min^51^, is inconsistent with a recycling model based on *in vitro* reaction conditions^50^. If polymerase recycling were a dominant mechanism for RNAPIII transcription, a substantially longer *in vivo* residence time to accommodate the additional rounds of transcription should be observed.

In contrast to RNAP-I and -III, RNAPII PIC components, also known as general transcription factors (GTFs) (Fig. 2a), exhibit substantially shorter residence times (Fig. 2b-d). We refined the previous estimates^38^ by applying an improved analytical approach, which includes incorporating endogenous tagged H2B as spike-in control (Extended Data Fig. 1b), utilizing a stretched exponential decay of H2B for correction (Extended Data Fig. 2), and analyzing only G1 phase cells to reduce potential cell-cycle heterogeneity (Extended Data Fig. 1a). Consistent with our earlier study^38^ and with recent competition-chromatin immunoprecipitation data^52^, GTFs generally have short residence times, ranging from 2-5 s as measured by SMT. In contrast, TBP^Spt15^ and TFIIA^Toa1^ show longer residence times of 28.5 s and 8.5 s respectively (Fig. 2c), as expected from their roles in stabilizing the early stages of hierarchical PIC assembly^53^. Although TFIIB was previously proposed to work in concert with TFIIA to stabilize the TBP-DNA interaction during the initial phase of PIC formation^54^, recent *in vitro* single-molecule imaging with purified human proteins showed unexpected dynamic binding behavior of TFIIB^55^, in line with our observation of a brief 2.1 s binding duration (Fig. 2c). Overall, we observe transient binding activity for the RNAPII PIC, contrasting with long-lived RNAP-I and -III PICs (Fig. 1d-k). This suggests that most RNAPII PICs may rapidly disassemble and reassemble on promoters for each transcription event^56,57^. However, the presence of a minor long-lived population of partial RNAPII PICs facilitating reinitiation^8, 52^, similar to the behavior of RNAP-I and -III PICs, cannot be excluded.

**Fig. 2.**
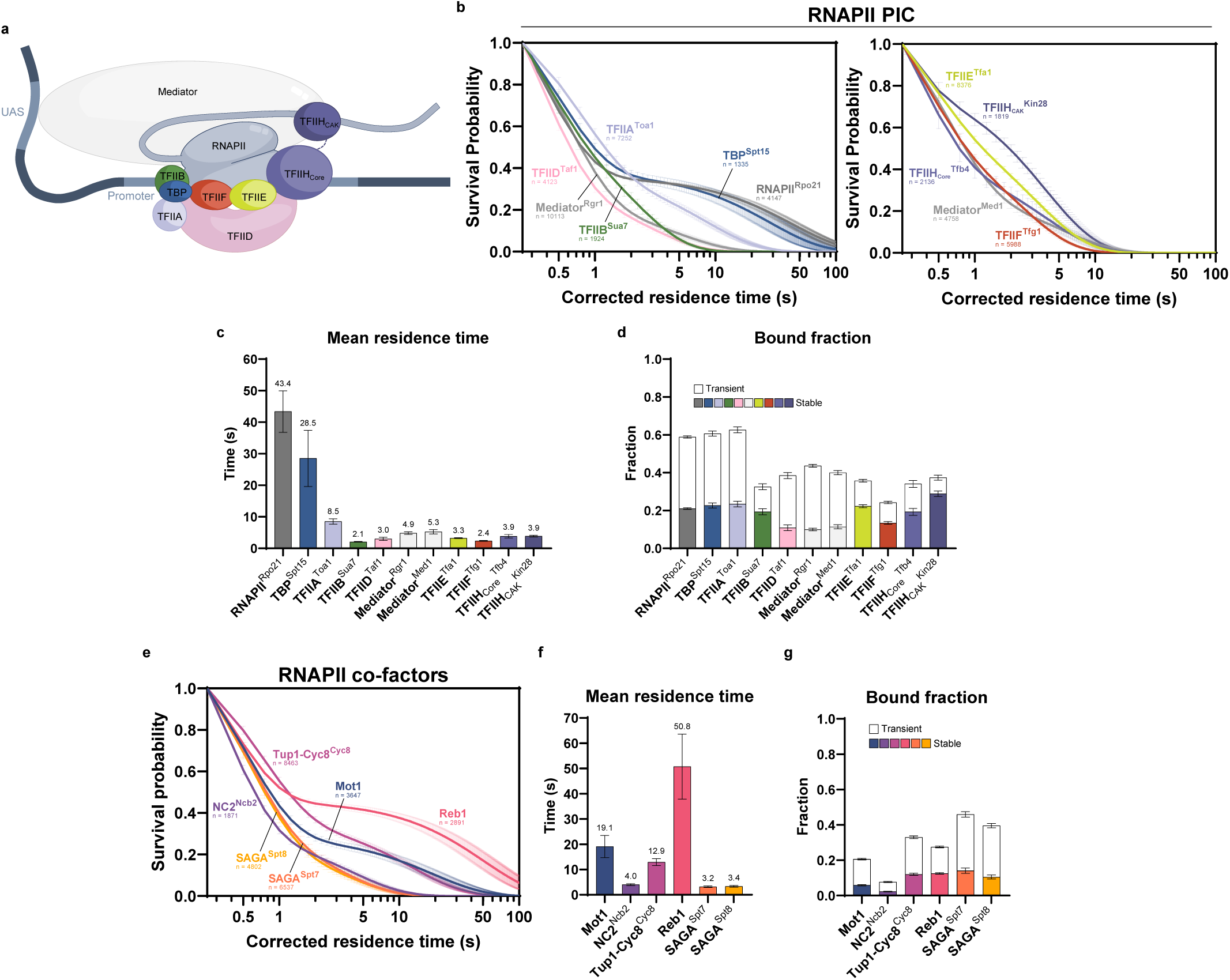
Short-lived RNAPII PIC with diverse dynamics of upstream co-factors. **a**, Schematic of RNAPII PIC. CAK = CDK activating kinase; UAS = upstream activation sequence. **b**, Survival probability of H2B-corrected residence times for RNAPII PIC (n = number of trajectories; mean value ± s.e.m.). **c**, Mean residence time of the stably bound fraction for RNAPII PIC (n = 5,000 resamplings; mean value ± s.e.m.). Given TBP involvement in all three nuclear RNAP PICs (Fig. 1d,h and 2a), we dissected its binding dynamics in different subnuclear compartments (Extended Data Fig. 3a,d-f); TBP^Spt15^ exhibits shorter residence time in the nucleoplasm (27.1 s) than in the nucleolus (40.5 s) (Extended Data Fig. 3a,d). This aligns with chromatin immunoprecipitation experiments demonstrating rapid TBP turnover at RNAPII promoters, moderate stability at RNAPIII promoters, and the slowest exchange at the RNAPI 35S promoter^79^. **d**, Transiently and stably bound fraction of RNAPII PIC (n = 5,000 resamplings; mean value ± s.e.m.). **e**, Survival probability of H2B-corrected residence times for upstream co-factors (n = number of trajectories; mean value ± s.e.m.). **f**, Mean residence time of the stably bound fraction for upstream co-factors (n = 5,000 resamplings; mean value ± s.e.m.). **g**, Transiently and stably bound fraction of upstream co-factors (n = 5,000 resamplings; mean value ± s.e.m.).

### Diverse binding dynamics of RNAPII upstream co-factors

We next explored the single-molecule dynamics of upstream co-factors in the RNAPII transcription pathway (Fig. 2e-g). These factors exhibit a wide range of chromatin-binding behaviors: TBP inhibitor Mot1 ATPase, co-repressor Tup1-Cyc8^Cyc8^, and general regulatory factor Reb1 (also a termination roadblock^14^) show longer residence times of tens of seconds (Fig. 2f). Reb1, in particular, exhibits a notable residence time of ∼50 s (Fig. 2f), highlighting the long-binding activity of its function as a nucleosome-displacement or pioneering’ factor as often observed across organisms^58–60^. The TBP inhibitor NC2^Ncb2^ and the SAGA^Spt7, Spt8^ complex involved in histone acetylation, bind for less than 4 s (Fig. 2f).

### Residence time analysis reveals stable and transient RNAPII elongation factors

Many RNAPII elongation-associated factors, previously assumed to be integral to the elongation complex based on gene-wide occupancies, display surprisingly diverse binding dynamics (Fig. 3a-g). Long binding times were observed for DSIF^Spt4^, Spt6, and Spn1, averaging ∼25 s (Fig. 3f), consistent with their roles as core components of the transcription elongation complex^61^. The cap-binding complex (CBC^Sto1^) displays a ∼20 s residence time (Fig. 3f), consistent with its function in protecting nascent RNA throughout elongation. In contrast, R-loop resolving factors TFIIS^Dst1^ and the THO complex^Hpr1^ exhibit dynamic binding of less than 2 s (Fig. 3f). Histone chaperone FACT^Spt16^ binds for ∼6 s. Capping enzymes (CE^Cet1^ and Abd1) and splicing factor U1 snRNP^Prp40, Prp39^ bind for ∼4-5 s, while Npl3, a multifaceted CTD and RNA-binding protein involved in RNA export and splicing, binds for ∼5 s (Fig. 3f). CTD modifiers (CTDK-1^Ctk1^, BUR kinase^Sgv1^, and Ess1) have residence times of ∼4-6 s, and histone methyltransferases (COMPASS^Set1^, Set2, and SET3 complex^Set3^) and histone deacetylase (Rpd3S^Rco1^) range from ∼3-7 s (Fig. 3f).

**Fig. 3.**
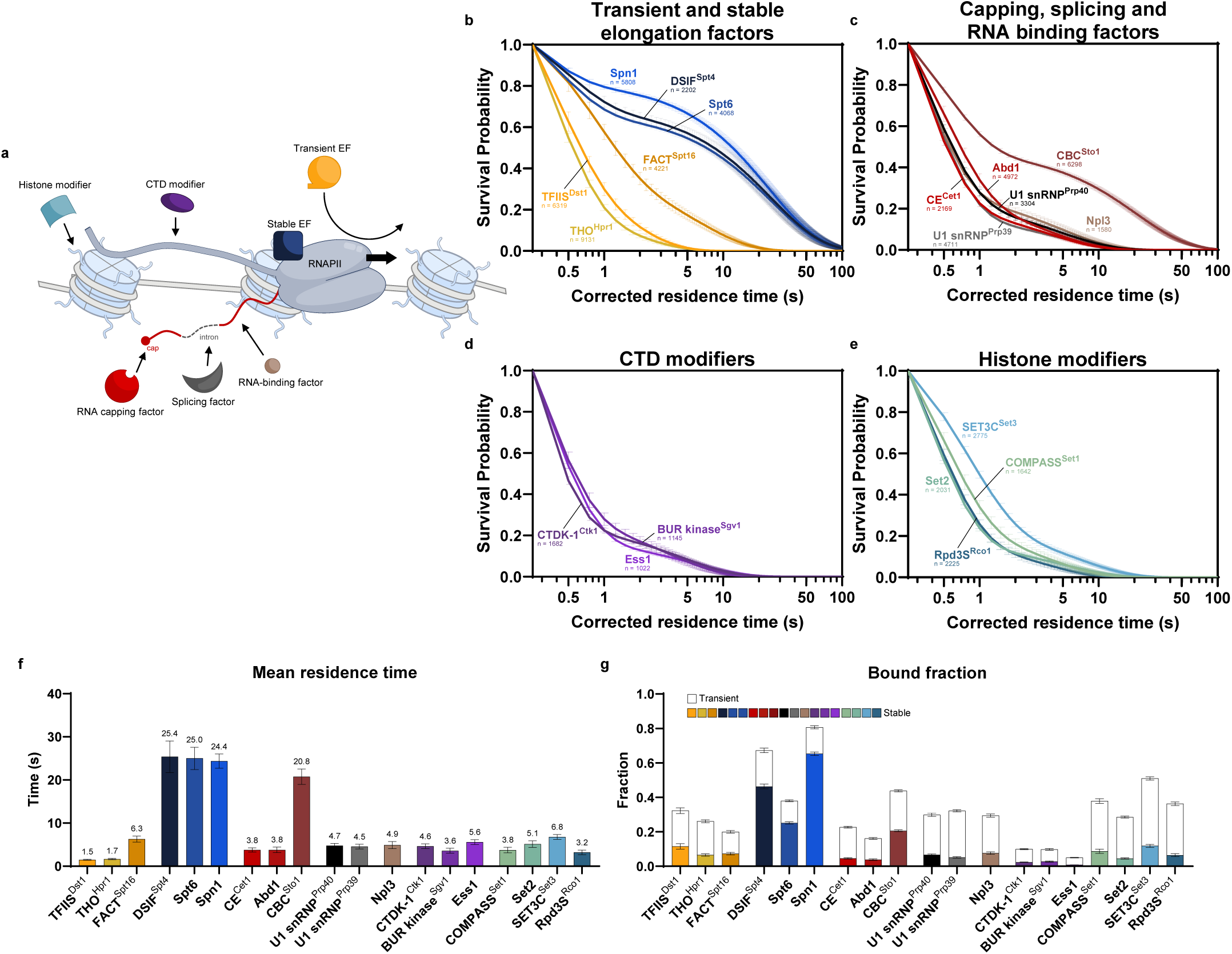
Unexpected transient binding of RNAPII elongation factors suggests dynamic interactions over constant association. **a**, Schematic of RNAPII elongation-associated factors. EF = elongation factor. **b-e**, Survival probability of H2B-corrected residence times for transient and stable elongation factors (b), capping, splicing and RNA binding factors (c), CTD modifiers (d), and histone modifiers (e) (n = number of trajectories; mean value ± s.e.m.). **f**, Mean residence time of the stably bound fraction for elongation-associated factors (n = 5,000 resamplings; mean value ± s.e.m.). DSIF was shown to be involved in both RNAPII and RNAPI elongation^80^; we observed DSIF^Spt4^ exhibits residence times of 24.4 s in the nucleoplasm and 8.1 s in the nucleolus, corresponding to its RNAPII- and RNAPI-associated activities, respectively (Extended Data Fig. 3b,d,f). The mechanistic role of DSIF^Spt4^ in elongating RNAPI remains unclear, and its unexpectedly transient dynamics within the nucleolus warrant further investigation. **g**, Transiently and stably bound fraction of elongation-associated factors (n = 5,000 resamplings; mean value ± s.e.m.).

Our findings suggest that many elongation factors interact transiently instead of persistently with elongating RNAPII. These global dynamics, likely dominated by highly expressed genes, provide insights into the elongation process: 1) gradual changes in the steady-state CTD phosphorylation pattern during elongation, described by genomic studies^62^, should be the consequence of multiple, transient enzyme-substrate interactions; 2) co-transcriptional capping^63^ and spliceosome recruitment^64^ should be highly efficient, occurring within a 5-s timescale; 3) co-transcriptional histone modification on the whole gene body likely requires repeated enzymatic interactions with RNAPII and histone tails, or multiple rounds of transcription; 4) the ∼25 s binding time observed for elongation factors DSIF^Spt4^, Spt6, and Spn1 should reflect the average duration for RNAPII elongation. For highly expressed genes of 1 kb average length (Extended Data Fig. 1d), we estimate the elongation speed of RNAPII to be 2.5 kb/min by SMT analysis, well within the 1-3 kb/min range documented on chromatin templates^65^.

### Contrasting kinetics of RNAPII termination factors and chromatin roadblocks

We expanded our analysis to factors involved in protein-coding gene termination (Fig. 4a-e), including CF1A^Pcf11^, CF1B^Hrp1^, CPF^Cft1, Ssu72^, and Rat1 complex^Rtt103^ (Fig. 4a), as well as the NNS complex^Nrd1^ for non-coding RNA gene termination^66^ (Fig. 4b). All examined factors display short residence times of ∼3 s (Fig. 4d), indicating transient recruitment and binding before dissociation. In contrast, Reb1, aforementioned as both a nucleosome-displacement factor^59^ and a termination roadblock^14^, has a long residence time of ∼50 s (Fig. 2f). Nsi1, a Reb1 homolog involved in RNAPI termination^67^, also exhibits a relatively stable residence time of ∼30 s (Extended Data Fig. 3c-e), suggesting its similar role as a roadblock for 35S gene termination. Overall, these findings underscore the striking differences in physiological timescales arising from factors involved in distinct termination mechanisms—either through dynamic interactions with the RNAPII enzyme and nascent RNA, or by functioning as stable chromatin roadblocks.

**Fig. 4.**
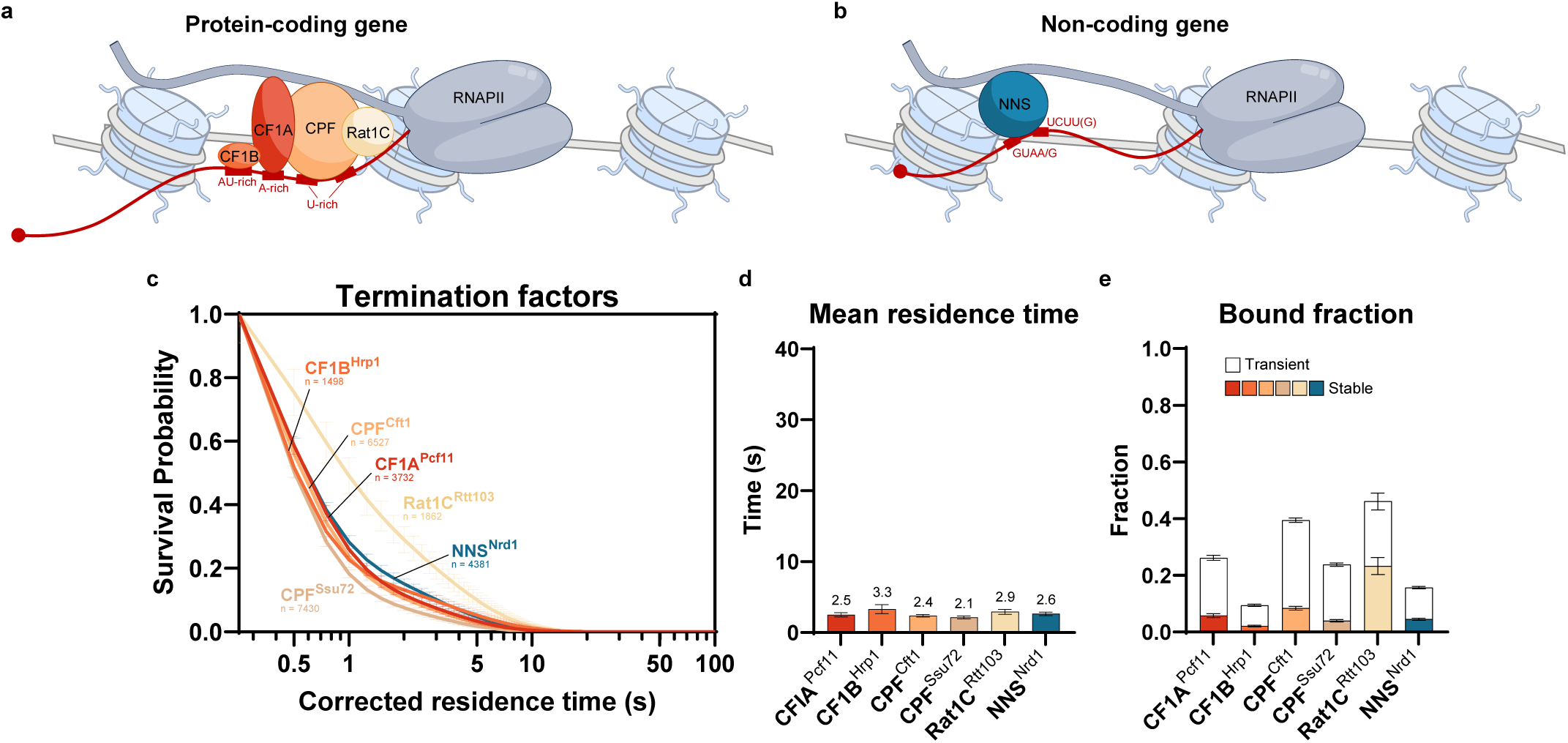
Brief engagement of RNAPII termination factors suggests rapid termination process. **a-b**, Schematic of RNAPII termination factors. **a**, For protein coding genes, the polyadenylation signal on nascent RNA consists of the AU-rich and A-rich elements upstream of the cleavage site, and the U-rich elements flanking it. These elements are bound by cleavage factor 1B (CF1B), cleavage factor 1A (CF1A), and the cleavage and polyadenylation factor (CPF), respectively. The Rat1 complex (Rat1C) contains RNA exonuclease activity to facilitate dissociation of polymerase from DNA^81^. **b**, Termination of non-coding RNAs is mediated by the NNS complex (Nrd1, Nab3, and Sen1), facilitated by binding to the Nrd1-binding motif GUAA/G and Nab3-binding motif UCUU(G) on the nascent transcript^82^. **c**, Survival probability of H2B-corrected residence times for termination factors (n = number of trajectories; mean value ± s.e.m.). **d**, Mean residence time of the stably bound fraction for termination factors (n = 5,000 resamplings; mean value ± s.e.m.). **e**, Transiently and stably bound fraction of termination factors (n = 5,000 resamplings; mean value ± s.e.m.).

### System-wide SMT of RNAPII machinery in a CTD truncation background

The RNAPII-CTD acts as a central hub for interactions with transcription-associated proteins throughout the transcription cycle^3–5^. Integrating and extending data from previous investigations^32^, we demonstrate that CTD truncation from 26 to 9 heptad repeats (CTD9) (Fig. 5a) impairs PIC formation, substantially reducing the stably bound fraction (binding frequency) of RNAPII^Rpo21^ and PIC components TFIIA^Toa1^, TFIIB^Sua7^, TFIIE^Tfa1^, TFIIF^Tfg1^, and TFIIH^Tfb4, Kin28^ (Extended Data Fig. 4). In contrast, upstream co-factors and early PIC components TBP^Spt15^ and TFIID^Taf1^ exhibit minimal changes in the stably bound fraction (Extended Data Fig. 4). As expected, many downstream interacting factors involved in stages beyond initiation show reduced stably bound fractions (Extended Data Fig. 4). Surprisingly, the residence time for the PIC and many other factors remains largely unchanged (Fig. 5b), indicating that the primary impact of reduced CTD length, at least for the PIC is on binding frequency (on-rate), rather than on binding stability (off-rate) as reflected by residence time. This observation aligns with a recent study in human cells, where RNAPII with only 5 CTD heptads associates with DNA less frequently, but once initiated remains pervasive for transcription^68^.

**Fig. 5.**
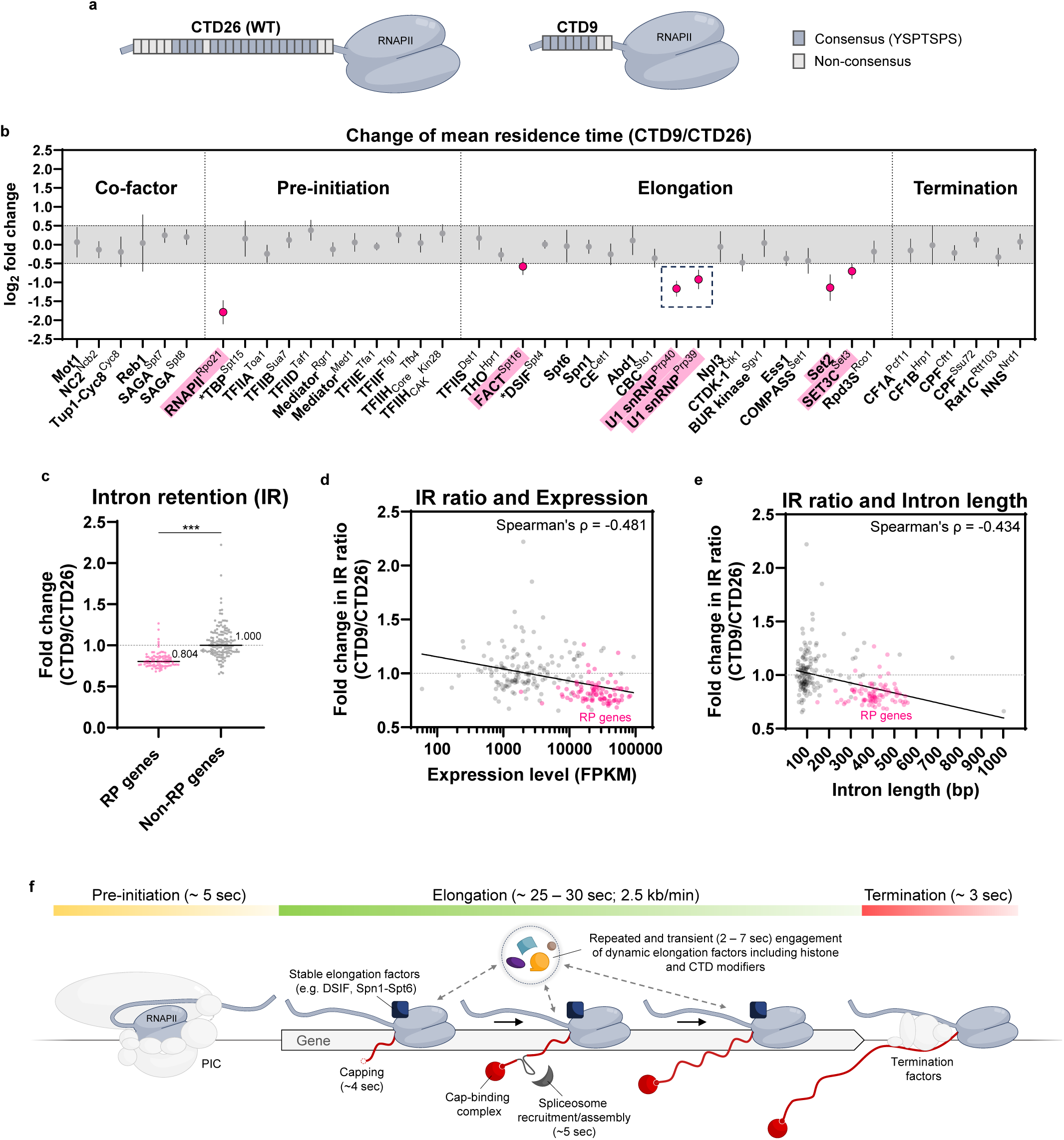
RNAPII-CTD truncation alters U1 snRNP binding dynamics and intron retention. **a**, Schematic of the disordered C-terminal domain (CTD) of RPB1, the largest subunit of RNAPII. WT CTD comprises 26 heptad repeats (CTD26) with the consensus sequence YSPTSPS. **b**, Log_2_ fold change in mean residence time of the stably bound population of RNAPII and associated proteins in CTD9 versus CTD26 background. Change greater than 0.5 or less than −0.5 is highlighted in pink (n = 5,000 resamplings; mean value ± s.e.m.). CTD truncation likely reduces RNAPII^Rpo21^ residence time due to compromised PIC recruitment and engagement with the Mediator^32^. For TBP^Spt15^ and DSIF^Spt4^ (marked with asterisks), only nucleoplasmic trajectories were analyzed, excluding nucleolar events to enrich for RNAPII-associated activities (Extended Data Fig. 3). **c**, Fold change in intron retention (IR) ratio for ribosomal protein (RP) genes and other intron-containing (Non-RP) genes (n = 4 biological replicates; median values were shown; Mann-Whitney Test, *** P = 2.01 x 10^-22^). **d-e**, Fold change in intron retention (IR) ratio against expression level (FPKM; Fragments Per Kilobase Million) (d) or intron length (e) for all intron-containing genes. n = 4 biological replicates. **f**, Model of RNAPII transcription cycle binding dynamics characterized by residence time analysis.

Intriguingly, a handful of elongation-associated factors, including U1 snRNP^Prp40, Prp39^, histone chaperone FACT^Spt16^, histone methyltransferase Set2, and histone deacetylase SET3 complex^Set3^, show sizeable reductions in residence time in the CTD9 mutant (Fig. 5b). Changes in the binding kinetics of histone chaperones and histone modifiers hint at possible epigenetic alterations from reduced transcription that invite further study. In addition, our finding that two subunits of the U1 snRNP, Prp40 and Prp39, show reduced residence times, suggesting a possible role of the CTD length in influencing co-transcriptional splicing (Fig. 5b). Among the spliceosome components, U1 snRNP is known for its direct interaction with the CTD, mainly through the subunit Prp40^69^. While 95% of yeast genes are intronless, two-thirds of ribosomal protein (RP) genes contain introns and are predominantly spliced co-transcriptionally^64^. To further investigate RNA splicing in the CTD9 mutant, we performed RNA-sequencing analysis, and found that the CTD9 truncation unexpectedly leads to a decrease of intron retention (IR), i.e. more mature mRNA molecules, specifically for RP genes, without changing their expression levels (Extended Data Fig. 5a). We observe a negative correlation between the IR ratio and gene expression level (Fig. 5d), or intron length (Fig. 5e), although this pattern is largely due to the unique properties of RP genes, which typically exhibit higher expression levels and longer introns than other intron-containing genes (Fig. 5d,e and Extended data Fig. 5b,c). Overall, our systematic SMT analysis of RNAPII machinery in the CTD9 background reveals altered U1 snRNP dynamics and suggests CTD length can influence RP gene pre-mRNA splicing.

## Discussion

In this work, we present a systematic quantitative analysis of dynamic chromatin interactions for RNAP-I, -II, and -III transcription machineries in living yeast, and perform system-wide analysis for the effect of CTD truncation on the dynamics of RNAPII-associated factors from initiation to elongation, RNA processing, and termination. We unexpectedly observed that splicing of RP genes is enhanced rather than diminished for the CTD9 mutant (Fig. 5c), suggesting a gain-of-function phenotype. In this regard, truncation of human RNAPII-CTD displays mixed behavior: shortening the CTD from 52 to 5 heptads results in splicing inhibition and lethality^70^, while a viable mutant with 25 heptads shows minimal effects on pre-mRNA splicing^26^, and another mutant with 26 heptads exhibits perturbation of alternative splicing^71^.

In yeast, a minimum of 8 CTD heptads is required for viability, but with substantial growth retardation^20^. Our study utilizes a CTD9 mutant (Fig. 5a), which maintains reasonable fitness^20^ for large-scale protein tagging and single-molecule tracking. Yeast RP genes are primarily spliced co-transcriptionally soon after the 3’ splice site is transcribed^64,72^. The known interaction between the CTD and U1 snRNP^69^ should facilitate the rapid search for splice sites during nascent RNA transcription and promote efficient co-transcriptional splicing^73^, yet splicing is not impaired but enhanced in the CTD9 mutant. It is conceivable that a truncated CTD might abnormally constrain the local search process for U1 snRNP, allowing it to locate splice sites even more efficiently, potentially accounting for the ∼2-fold reduction in U1 snRNP residence time (Fig. 5b), and the enhancement of RP gene pre-mRNA splicing (Fig. 5c). Additional considerations, such as altered phosphorylation on truncated CTD heptads affecting U1 snRNP activity and RP gene-specific adaptation responses, are possible. Changes in RNAPII elongation speed might also influence splicing efficiency^72^, but this does not seem to be altered in the CTD9 mutant, as stable elongation factors DSIF^Spt4^, Spt6, and Spn1 exhibit residence times comparable to wild type (WT) (Fig. 5b). Further mechanistic insights will require knowledge of CTD9 modification state, and development of *in vitro* single-molecule assays incorporating the RNAPII-CTD to study co-transcriptional splicing.

PICs of RNAP-I and -III are characterized by prolonged chromatin engagement, with the most stable components ranging from tens to hundreds of seconds (Fig. 1f,j), in line with their roles in maintaining a default ON’ state for rRNA and tRNA gene transcription. In contrast, most RNAPII PIC components excepting TBP are marked by strikingly brief chromatin binding of several seconds (Fig. 5f), reflecting flexibility for rapid, inducible regulation of mRNA and non-coding RNA expression. Our supplementary kinetic modeling of temporal occupancy, defined as the percentage of time a chromatin target is occupied by a factor, indicates that target sites of long-binding RNAPI PIC components CF^Rrn6^ and UAF^Rrn9^, and RNAPIII PIC component TFIIIB^Bdp1^ exhibit occupancy close to 100% (Extended Data Fig. 6). This supports the model where RNAPI and RNAPIII promoters are constantly bound by these PIC components respectively and are constitutively accessible. In contrast, chromatin target sites associated with RNAPII PIC components have a much lower occupancy of ∼10%, as suggested by previous study^38^.

The prolonged binding of the RNAPI PIC components CF and UAF observed in live yeast (Fig. 1f) differs from *in vitro* experiments using yeast whole-cell extracts, where CF dissociates upon transcription initiation while UAF remains bound^36^. However, experiments with HeLa cell nuclear extracts show that the human homologs of CF (SL1) and UAF (UBF) continue to stay bound to the promoter during multiple rounds of transcription^37^. The basis of these differences is not known and requires further *in vitro* studies to be reconciled with the observations in living cells.

Our dynamic model of elongation-associated factor binding (Fig. 5f) could explain how the RNAPII-CTD enables numerous interactions, linking histone modification, CTD modification, and RNA processing to transcription elongation. Transient association would allow increased CTD space for a greater number of CTD-interacting macromolecular complexes over time without steric hindrance, especially as effective CTD binding typically requires two consecutive heptad repeats^74^. Furthermore, transient interactions may enrich the repertoire of biological controls over elongation rates and co-transcriptional activities, enabling more adaptable regulation compared to a continuously bound set of elongation factors.

The binding duration of transcription termination factors *in vivo* is surprisingly brief, averaging only ∼3 s (Fig. 5f). This contrasts with an early model of termination for the long *MDN1* gene, based on fluctuation analysis of fluorescent nascent RNA in live yeast, which suggested a termination time of approximately 70 s^75^. The rapid turnover implied by our measurements suggests that assembly and disassembly of termination complexes should occur much faster than previously considered. The different findings could be due to *MDN1* gene-specific effects compared to the global average, or to the nascent RNA reporter lingering on-site longer than the actual termination event. While several binding events of ∼3 s before effective termination remains possible, repetitive binding should be accommodated within the 40-s average residence time of RNAPII on a 1 kb yeast gene—encompassing initiation, elongation, and termination (Fig. 5f).

The transcription protein dynamics revealed in our live-cell study likely reflect activities that are predominantly for highly expressed genes. While all these factors are crucial in transcription and exhibit considerable occupancy on transcribed genes^76^, it is important to note that they may also be involved in other cellular functions beyond transcription, such as cell cycle regulation and DNA repair. In the future, it would be of interest to measure single-molecule dynamics within specific gene loci *in vivo*^41,77,78^ and employ *in vitro* single-molecule imaging on native or engineered gene templates^9^ to elucidate targeted gene interactions by the transcription machinery.

## Acknowledgments

We thank Luke Lavis for providing Janelia Fluor dyes; Chuofan Yu and Theresa Mai for their preliminary contributions to the SMT dataset; Craig Kaplan and Jeffry Cordon for helpful discussions; Members of the Wu laboratory for comments. This study was supported by National Institute of Health grant GM132290 and R35GM149291 (C.W.) and the Croucher Foundation (Y.H.L.).

## Author Contributions Statement

Y.H.L. designed the experiments. Y.H.L. performed the experiments and analyzed the data. C.L. and S.W. assisted in experiments. Y.H.L., and C.W. wrote the manuscript. C.L., S.W. contributed equally.

## Competing Interests Statement

The authors declare no competing interests.

## Statistics and reproducibility

Statistical analyses were performed using GraphPad Prism (v9.4.0) or RStudio (v2021.09.1+372; R (v4.1.2)). Values are mean ± s.e.m. unless otherwise stated. Statistical tests and P values are reported in figure legends. For reliable fitting and statistical evaluation, data obtained from at least two independent imaging sessions were resampled by bootstrapping 5,000x. Representative microscopic images from at least two independent acquisitions are shown in the figures.

## Supplementary information

### Methods

**Extended Data Fig. 1-7**

**Supplementary Table 1. Yeast strains for this study.**

**Supplementary Video 1. Example video of fast tracking.** Fast tracking of RNAPII^Rpo21^, RNAPI^Rpa190^, and RNAPIII^Ret1^. Nucleolus and nuclear envelope outlined in solid white, plasma membrane in dashed white. Asterisks indicate nucleoli. Scale bar: 1.0 µm. Recorded frame rate: 10 ms per frame. Playback speed: 20 ms per frame.

**Supplementary Video 2. Example video of slow tracking.** Slow tracking of RNAPII^Rpo21^ with H2B^Htb1^ spike-in control. Nucleolus and nuclear envelope outlined in solid white, plasma membrane in dashed white. Asterisks indicate nucleoli. Scale bar: 1.0 µm. Recorded frame rate: 250 ms per frame. Playback speed: 100 ms per frame.

## Methods

### Yeast construction

All *Saccharomyces cerevisiae* strains in this study (Supplementary Table 1 and Extended Data Fig. 7) are isogenic derivatives of BY4741 with *pdr5* deletion for enhanced HaloTag ligand labeling, and with GFP fused to endogenous loci of ER (Elo3) and nucleolar (Gar1) markers. Strains with *pdr5*Δ and GFP markers exhibit growth identical to the parental BY4741 strain^32^. The CTD9 mutation was engineered via knock-in at the endogenous locus and retains the tip domain and non-consensus heptads^32^.

### Microscope setup

The microscope setup has been previously described^32^. Briefly, imaging was conducted using a custom-built Zeiss Axio Observer Z1 microscope (Zeiss, Germany) with a 150X glycerin immersion objective (Zeiss, Germany). Data were recorded using an EM-CCD camera (C9100-13, Hamamatsu Photonics, Japan) with a 16 µm physical pixel size and a 107 nm pixel size in recorded images, controlled by HCImage (v5.0.1). Laser excitation wavelengths were 488 nm for GFP, 555 nm for JF552, and 637 nm for miRFP670nano3. Laser power and alignment were routinely measured and adjusted throughout imaging sessions to ensure consistency.

### Single-molecule tracking

Yeast cultures in synthetic complete (SC) medium were incubated with JF552-HaloTag ligand during early logarithmic growth (OD_600_ = 0.2-0.3) for 4 hours. HaloTag fusion proteins were labeled with adjusted dye concentrations to ensure sparse labeling. After at least six washes, cells were attached to concanavalin A-coated coverslips in Attofluor™ cell chambers (Invitrogen, USA) and imaged at room temperature. Initial illumination with a 555 nm laser produced a strong nuclear signal, followed by a transient shift of fluorophores into a metastable dark state. JF552 spontaneously and randomly returned to a fluorescent state, allowing for sparse single-molecule detection. G1 cells were selected for analysis using GFP markers^32^ (Extended Data Fig. 1a). Each imaging session lasted up to two hours, equivalent to one cell cycle in SC at room temperature, typically covering 10-40 fields of view. For robust statistics, all datasets included at least 1,000 trajectories from at least two independent imaging sessions. Laser exposure during imaging showed no substantial effect on cell cycle progression, as previously demonstrated^32^. Single-molecule localization and trajectory linking were performed using Diatrack (v3.05)^83^, with a maximum jump set at 6 pixels (642 nm) for fast tracking and 3 pixels (321 nm) for slow tracking^32^. Nuclear masks were manually drawn to exclude cytoplasmic localizations. Single-step disappearances of bound molecules, observed in both fast and slow tracking from our previous study, suggest accurate identification of single-molecule signals^32^.

### Fast tracking

Fast tracking was conducted at 10 ms/frame with continuous 555 nm laser irradiation at ∼1 kW/cm^2^ for analyzing bound and free molecule dynamics. Static localization error was ∼20 nm^32^. To minimize the inclusion of cytoplasmic diffusing trajectories, we exclusively selected G1 cells exhibiting minimal nuclear drift during imaging^32^. Transitioning trajectories identified by vbSPT^84^ were separated into single-state segments. Each trajectory was then classified as chromatin-bound for the slow population or freely diffusing for the fast population based on diffusivity, geometry, and angular orientation using a multi-parameter approach described previously^32^.

We functionally attribute the slow trajectories to chromatin binding, either directly or indirectly, as evidenced by their enrichment in subnuclear compartments with specific binding sites^32^ (Fig. 1c). These trajectories exhibit a mean apparent diffusion coefficient similar to that of histone H2B, yet they are typically an order of magnitude slower than the fast, freely diffusing population^32^ (Extended Data Fig. 7). Notably, this chromatin-associated population was absent in the HaloTag control experiments^32^. Biological perturbations affecting factor binding typically result in a reduced fraction of this population^32,38,39^.

We acknowledge that protein factors may exist independently or as part of various large complexes, resulting in freely diffusing trajectories with intermediate apparent diffusion coefficients. As this study primarily focuses on the bound population, we adhere to a simplified 2-state model to classify the principal states as free and bound.

### Slow tracking

Slow tracking was conducted at 250 ms/frame using continuous 555 nm laser irradiation at a reduced power of ∼0.035 kW/cm^2^ to minimize photobleaching and improve residence time resolution. To account for blinking or transient defocalization of bound molecules, we allowed gaps of up to one frame between localizations and linked them as a single trajectory if they were within 3 pixels (321 nm)^32^. We spiked the imaging culture with Halo-H2B cells labeled with the nuclear marker Pus1-miRFP670nano3 and co-imaged the two strains in the same field of view^32^ (Extended Data Fig. 1b). A survival curve (1-CDF) against time (t) was plotted and fitted with a two-exponential decay model, incorporating a stretched exponential decay of H2B to account for experimental artifacts such as photobleaching, nuclear and chromatin movement, microscope drift, and dye photophysics (2-exp_StrCORR_; Extended Data Fig. 2d):

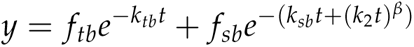

In this model, *k_tb_*and *k_sb_* are the dissociation constants for the transient and stable binding populations, respectively. *k_2_* is the dissociation constant for stable binding population of H2B fitted with a stretched exponential decay, and *β* is the stretching exponent ranges between 0 and 1. The fractions of the two components, *f_tb_* and *f_sb_*, add up to 1 (Extended Data Fig. 2d). The transient population, with sub-second residence times, is minimally affected by experimental artifacts; thus its dissociation rate is not corrected with H2B decay to avoid over-fitting. The stably bound fraction was determined by multiplying the total bound fraction from fast tracking with the stably bound fraction (*f_sb_*) from slow tracking. One limitation of this approach is the differing exposure times used for fast and slow tracking (10 ms/frame versus 250 ms/frame), as the longer frame rate in slow tracking may not effectively capture highly transient binding events. However, the estimated stably bound fraction still provides a valuable metric for assessing changes in binding kinetics. Stable binding is interpreted as a proxy for protein activity at specific sites^32,38–41,85^. However, we regard transient binding as a neutral description of duration in two-component fitting, rather than as merely non-specific site binding^32^.

### Residence time correction with stretched exponential decay of H2B

The survival curve of chromatin-binding proteins is typically fitted with a two-exponential decay model as stable and transient populations (Extended Data Fig. 2k,l). The residence time of the stable population is corrected for experimental artifacts like photobleaching, nuclear and chromatin movement, microscopic drift, and dye photophysics^32^, using the slow decay constant from a separate two-exponential decay fit of H2B (2-exp_ExpCORR_) (Extended Data Fig. 2a-d). For optimal accuracy, the Halo-H2B sample should be imaged on the same day^85^ or added as a spike-in control^32^ (Extended Data Fig. 1b). Studies in mammalian cells suggest that a power-law model may more accurately represent the chromatin binding behavior of transcription factors^86,87^. This model proposes that transcription factors bind across a continuum of affinities at multiple sites and cannot be distinctly categorized into transient and stable populations; thus, specific residence times cannot be derived. In this approach, the survival curve of the protein of interest is first adjusted using the slowest exponential decay constant from a three-exponential decay fit of H2B, followed by fitting to a power-law distribution^86,87^.

Given that classic kinetic parameters like rate constant and residence time are not obtainable with power-law fitting^88^, we explored an alternative model to account for the continuum-like feature of the survival curve, using the three nuclear RNAPs in yeast as test cases. Similar to findings with mammalian transcription factors, we have observed that a two-component model with standard H2B correction (2-exp_ExpCORR_) does not accurately capture the decay curve for the three RNAPs (Extended Data Fig. 2e-h), similar to previous findings by others on yeast RNAPII^89^. We questioned whether refining the modeling of the H2B decay for correction could enhance the curve fitting of the RNAPs. Our analysis revealed that fitting the H2B decay using a combination of exponential and stretched exponential decay (1e1s) is superior to a simple two-exponential decay (2-exp) (Extended Data Fig. 2b-d). This revised model describes the experimentally observed bound H2B as two populations: a transient population described by standard exponential decay, and a stable population characterized by a complex stretched decay’ profile, which accounts for multiple experimental artifacts. Stretched decay has also been adapted for modeling fluorescence decay in complex systems^90,91^. Notably, applying the stretched decay of H2B for correction (2-exp_StrCORR_) refines the two-exponential fitting of the three nuclear RNAPs in yeast, with marked improvement on residence time estimation and resolution, particularly for RNAPI and RNAPII, which have longer residence times (Extended Data Fig. 2d-j). Further research is needed to evaluate the applicability of this approach in organisms beyond yeast, especially considering the faster chromatin dynamics observed in yeast compared to mammalian cells. For instance, the apparent diffusion coefficient of bound H2B captured by SMT at 10 ms/frame is ∼0.05 µm^2^/s in yeast^32^ and ∼0.01 µm^2^/s in mammalian cells^92^.

### RNA-Sequencing

Yeast cells grown in SC medium were harvested and homogenized with bead beating. Total RNA was isolated using the SPLIT RNA Extraction Kit (Lexogen, Austria) and treated with DNase I (NEB, USA) to eliminate genomic DNA. rRNA was depleted using the Ribocop for Yeast kit (Lexogen, Austria), and sequencing libraries were prepared with the CORALL Total RNA-Seq V2 kit (Lexogen, Austria). Libraries were sequenced on the Element AVITI™ System (Element Biosciences, USA) (paired-end 2×75 bp). Four biological replicates were performed, yielding approximately 100M reads each. Adapter sequences were trimmed using Cutadapt (v1.18), and reads were aligned to the *S. cerevisiae* genome (Genome Assembly R64) using STAR (v2.6.1a). PCR duplicates were removed by collapsing reads with identical mapping coordinates and UMI (unique molecular identifier) sequences. Expression level was calculated using Mix^2^ (v1.4.0.2; Lexogen, Austria). Intron retention (IR) ratio was quantified using IRFinder-S (v2.0)^93^.

---

**Extended Data Fig. 1.**
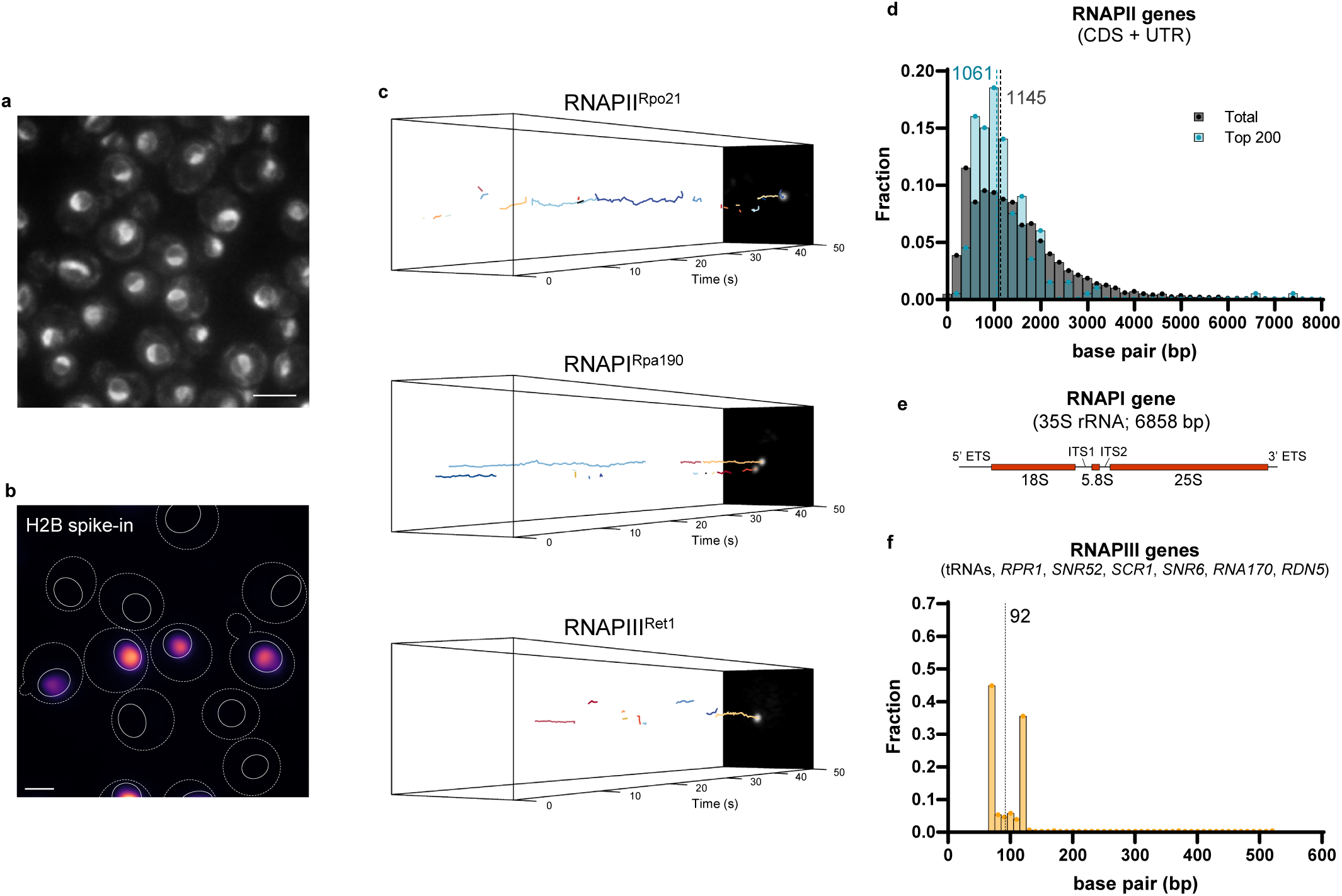
Single-molecule tracking of eukaryotic nuclear RNA polymerases. **a**, Strain with ER (Elo3) and nucleolar (Gar1) markers fused with GFP for cell cycle identification^32^. Scale bar: 4.0 µm. **b**, Halo-H2B spike-in control identified with Pus1-miRFP670nano3 nuclear marker. Cell membrane indicated by dashed line; nuclear envelope by solid line. Scale bar: 2.0 µm. **c**, Kymograph of slow tracking trajectories for the three nuclear RNAPs. **d-f**, Geometric mean of RNAP-II (d), -I (e), and -III (f) genes. **d**, Length of RNAPII gene coding sequence (CDS) with untranslated region (UTR) obtained from Tuller et al., 2009^94^. Top 200 mRNA genes for yeast in SC medium obtained from Miura et al., 2008^95^. **e**, ETS = external transcribed spacer; ITS = internal transcribed spacer. **f**, 275 tRNA genes, *RPR1*, *SNR52*, *SCR1*, *SNR6*, *RNA170*, and 150 copies of 5S rRNA gene (*RDN5*) with a length of 120 bp were set for calculation^96,97^.

**Extended Data Fig. 2.**
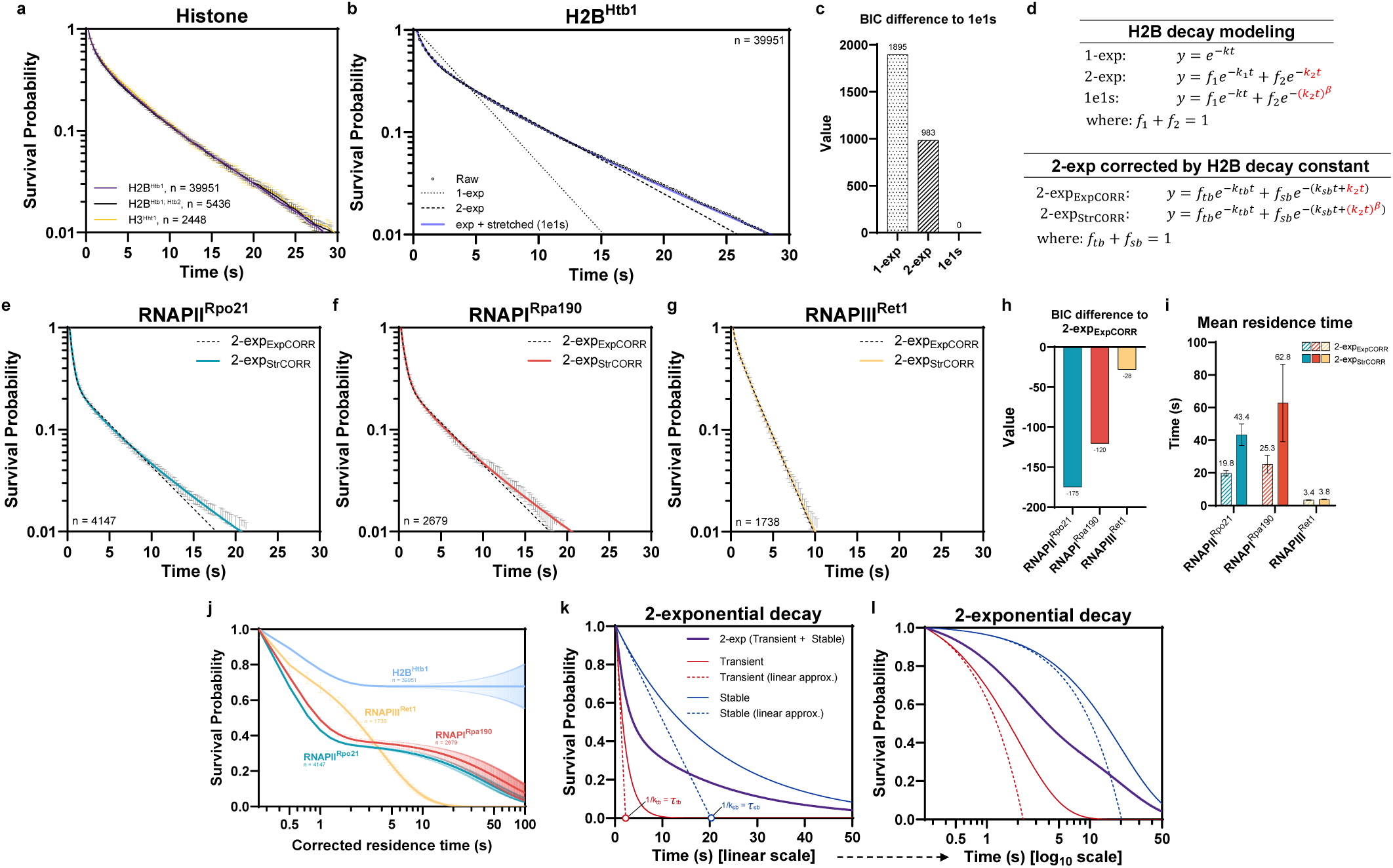
Improved residence time correction using stretched exponential decay of H2B. **a**, Labeling one copy of H2B (Htb1), both copies of H2B (Htb1 and Htb2), or one copy of H3 (Hht1) resulted in essentially identical survival probabilities (n = number of trajectories; mean value ± s.d.). **b**, Fitting of Halo-H2B survival probability plot with one-exponential decay (1-exp), two-exponential decay (2-exp), and one exponential plus one stretched exponential decay (1e1s) (n = number of trajectories). **c**, BIC difference of fittings compared to 1e1s. **d**, Equations of H2B decay modeling for residence time correction. *f* = fraction; *k* = decay constant; *β* = stretching exponent. **e-g**, Survival probability plot of RNAP-II (e), -I (f), and -III (g) fitted with two-exponential decay, and corrected by either a standard exponential decay (2-exp_ExpCORR_) or a stretched exponential decay (2-exp_StrCORR_) from stable population of H2B (n = number of trajectories; mean value ± s.d.). **h**, BIC difference for 2-exp_ExpCORR_ versus 2-exp_StrCORR_. **i**, Mean residence time of nuclear RNAPs calculated using either 2-exp_ExpCORR_ or 2-exp_StrCORR_ (n = 5,000 resamplings; mean value ± s.e.m.). **j**, Survival probability of H2B-corrected residence times for H2B, RNAP-II, -I, and -III (n = number of trajectories; mean value ± s.e.m.). **k**, Mean residence time of exponential decay (*τ*) can be estimated graphically through linear approximation (approx.) by taking the slope of the tangent at the initial point, with the x-intercept representing 1/k. **l**, The x-axis (time) of the survival probability plot on a log_10_ scale enhanced visualization of the transient and stable populations. Figure legend as in k.

**Extended Data Fig. 3.**
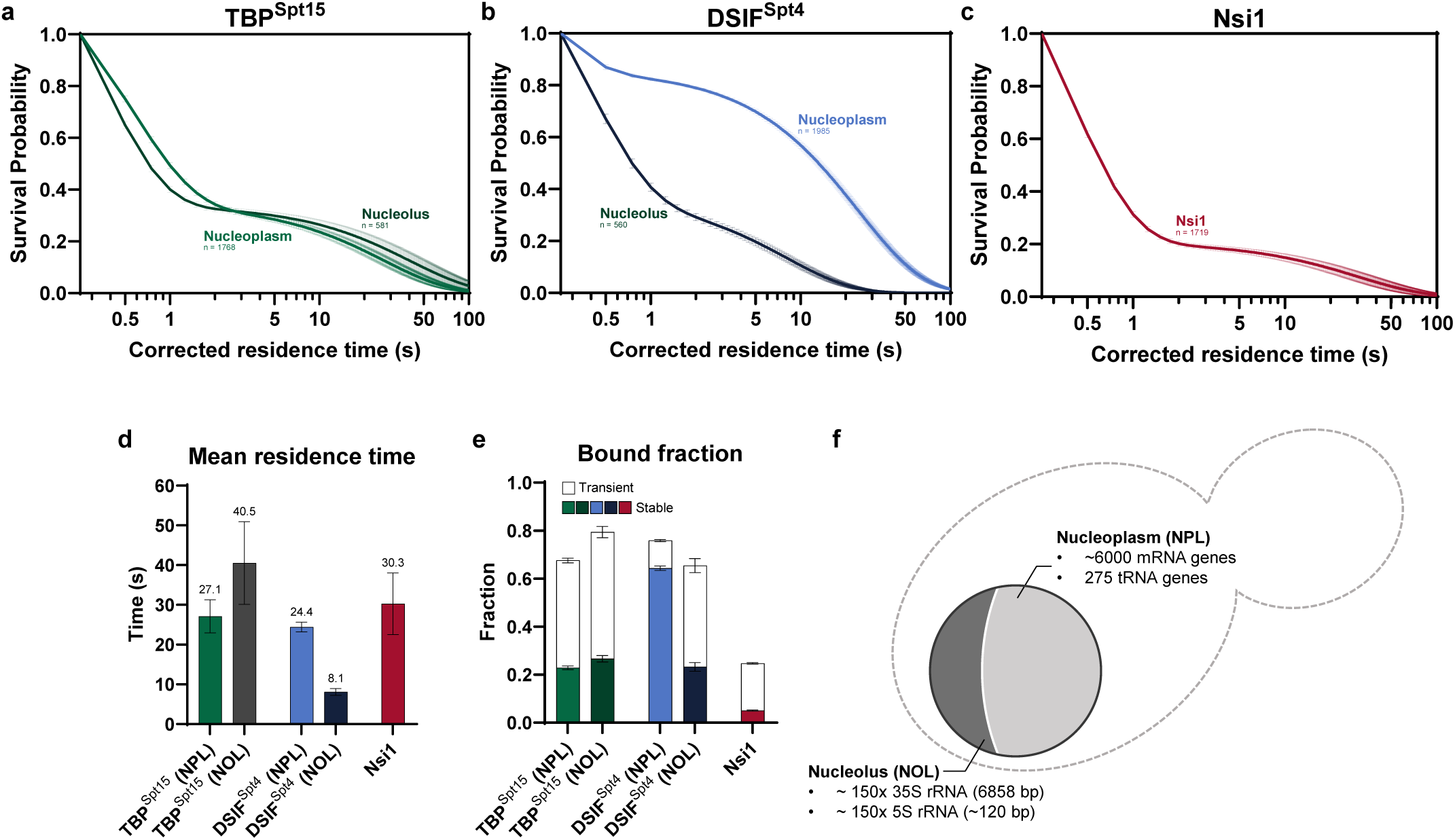
RNAPI-associated dynamics for TBP^Spt15^, DSIF^Spt4^, and Nsi1. **a-c**, Survival probability of H2B-corrected residence times for TBP^Spt15^ (a), DSIF^Spt4^ (b) in nucleoplasm (NPL) and nucleolus (NOL), and Nsi1 (c) (n = number of trajectories; mean value ± s.e.m.). **d**, Mean residence time of the stably bound fraction (n = 5,000 resamplings; mean value ± s.e.m.). **e**, Transiently and stably bound fraction (n = 5,000 resamplings; mean value ± s.e.m.). **f**, Spatial distribution of 35S rRNA, 5S rRNA, tRNA genes, and mRNA genes in nucleoplasm and nucleolus.

**Extended Data Fig. 4.**
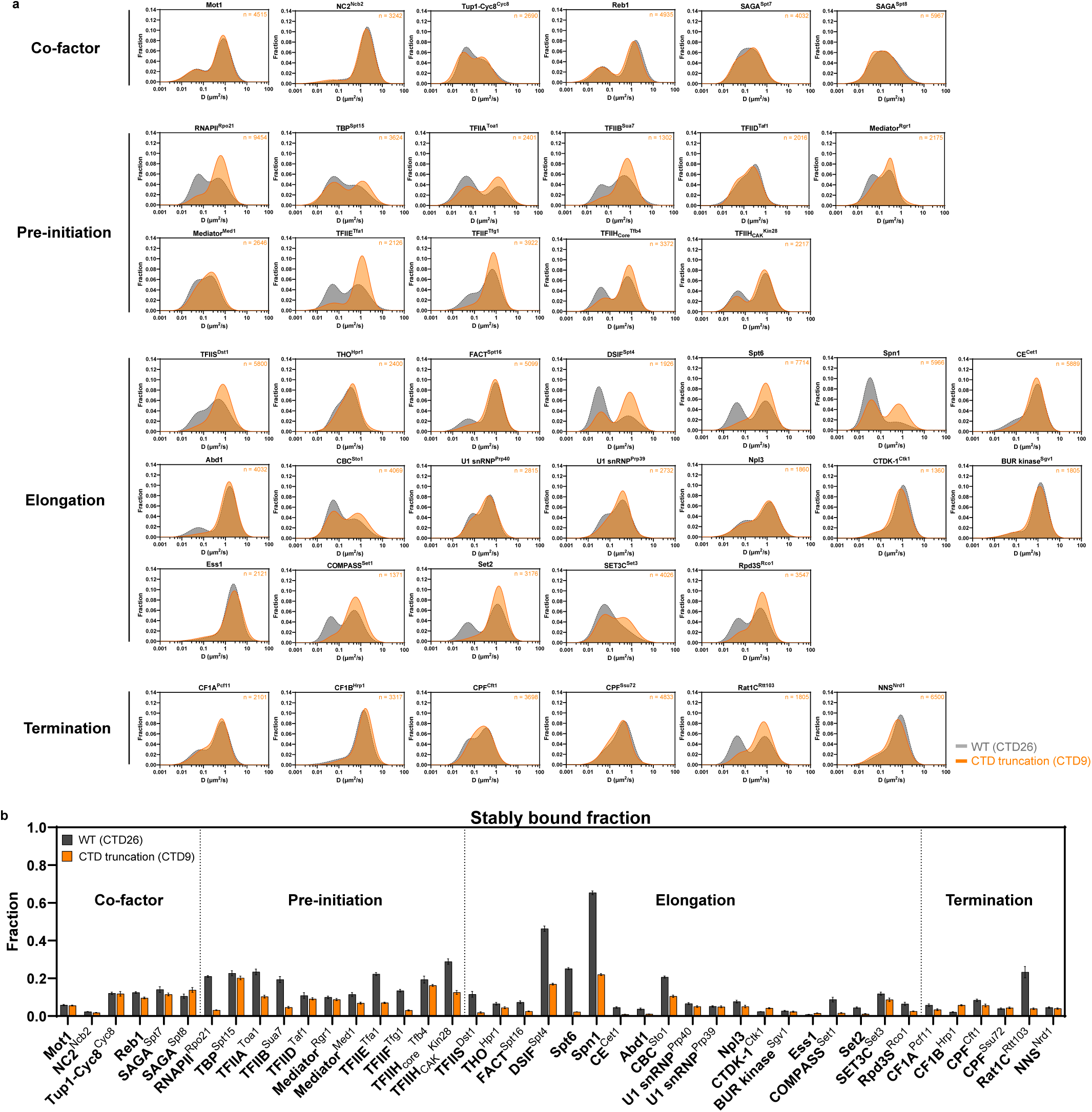
Single-molecule dynamics of RNAPII and associated factors in WT and CTD9 mutant. **a**, Two-component Gaussian fit of apparent diffusion coefficient (D) histograms (n = number of trajectories for CTD9 mutant; number of trajectories for WT in Extended Data Fig. 7). **b**, Stably bound fraction of RNAPII and associated factors in WT and CTD9 mutant (n = 5,000 resamplings; mean value ± s.e.m.). While CTD truncation impairs PIC formation, as observed in the figure and in our previous study^32^, changes in chromatin-binding fraction for some factors downstream of initiation (e.g. CTDK-1^Ctk1^ and CF1B^Hrp1^) are uncorrelated with the anticipated reduction in transcription. These varied responses to CTD truncation might reflect underlying cellular adaptations or complex regulatory mechanisms that are not yet fully understood, and will require further targeted analysis.

**Extended Data Fig. 5.**
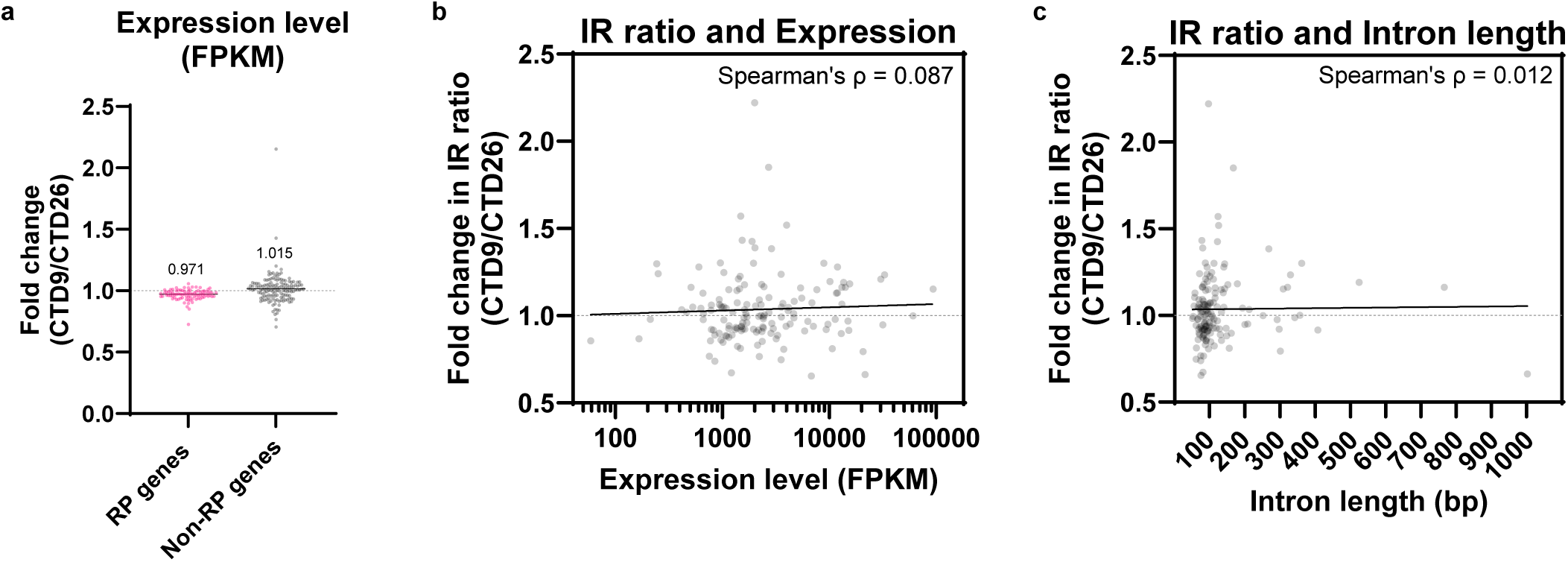
Expression level and intron retention ratio of intron-containing genes in WT and CTD9 mutant. **a**, Fold change in expression level (FPKM; Fragments Per Kilobase Million) for ribosomal protein (RP) genes and other intron-containing (Non-RP) genes (n = 4 biological replicates; median values were shown). **b-c**, Fold change in intron retention (IR) ratio against expression level (FPKM; Fragments Per Kilobase Million) (b) or intron length (c) for intron-containing non-RP genes (n = 4 biological replicates).

**Extended Data Fig. 6.**
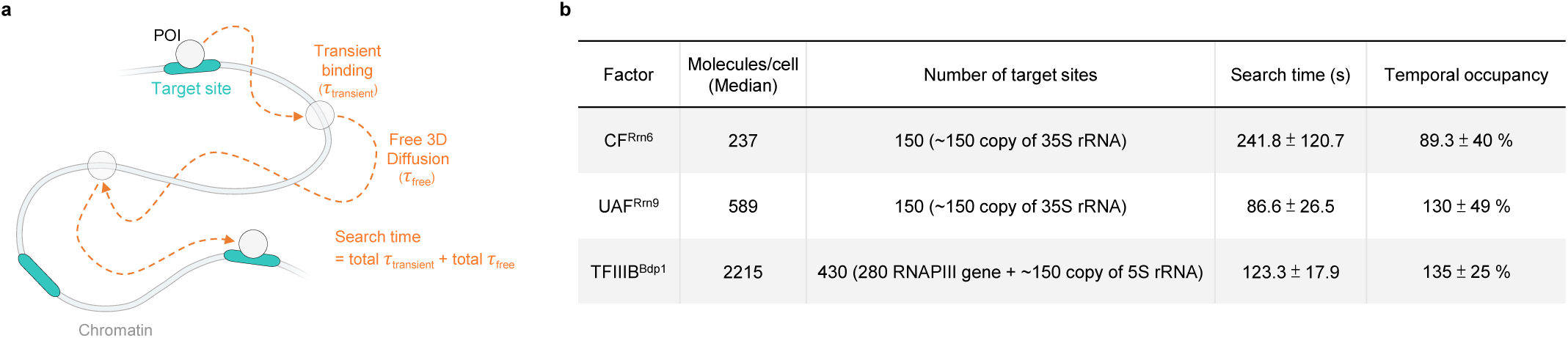
High temporal occupancy by PIC components suggests persistent accessibility of RNAPI and RNAPIII promoters. **a**, Simplified model for the diffusive search of a protein factor between two specific chromatin target sites in the nucleus. POI = protein of interest. **b**, Temporal occupancy measures the percentage of time a chromatin target is occupied by a factor, while search time calculates the average duration a protein molecule spends between stable binding events (mean value ± s.e.m.). The model integrates data on binding dynamics from fast and slow tracking, along with the estimated number of molecules^98^ and target sites per cell^96,97^. Detailed calculations are available in prior studies^38,58^. It is important to note several limitations: slow tracking at 250 ms/frame may not capture exceedingly brief binding events reported by fast tracking at 10 ms/frame; molecule counts from Ho et al., 2018^98^, are estimates from multiple datasets and may differ in our conditions; for simplicity and consistency with the SMT data, only the molecule count of one subunit of a complex is used; target sites are based on estimated counts for known RNAPI and RNAPIII genes^96,97^, potentially missing unconventional binding sites; additionally, factors might sample more frequently at sites on gene promoters with higher firing rates. Despite these considerations, this modeling approach provides valuable insights but should be viewed as supplementary and interpreted with caution.

**Extended Data Fig. 7.**
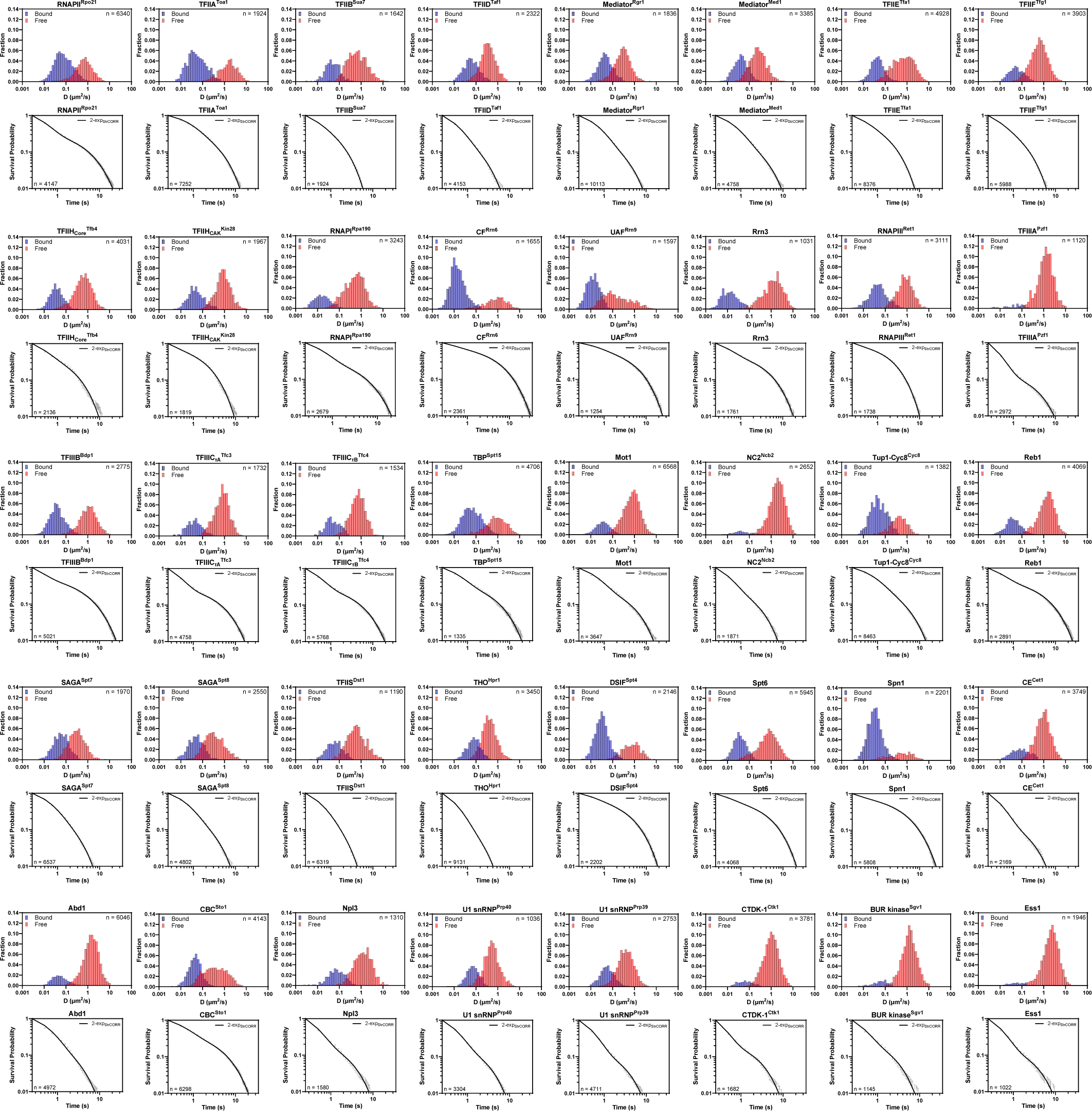
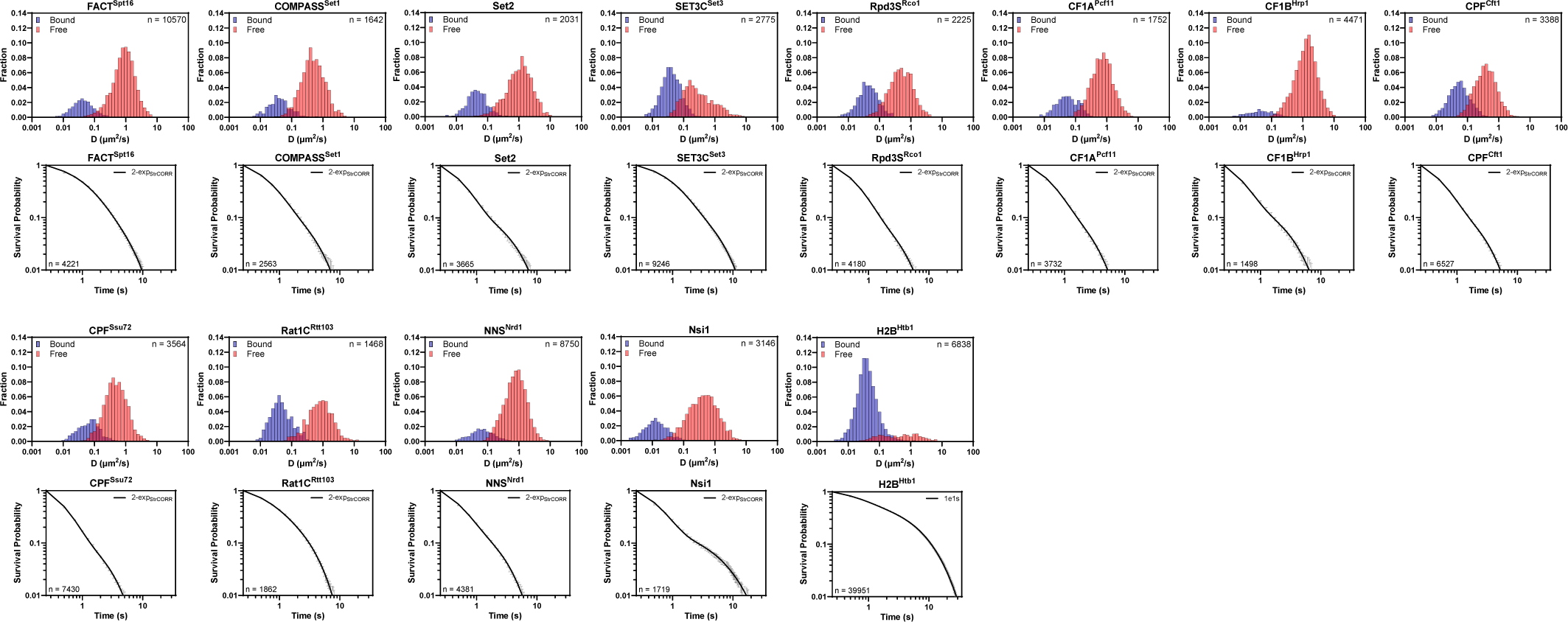
Apparent diffusion coefficient histograms and survival probability of factors examined. Superscripts indicate the tagged representative subunits of complexes or yeast-specific nomenclature. For survival probability (1-CDF) plots, lines indicate 2-exp_StrCORR_ fit and error bars indicate s.d. of raw values. Fast tracking data (apparent diffusion coefficient histograms) of RNAPII PIC components, SAGA^Spt7^, ^Spt8^, RNAPI^Rpa190^, RNAPIII^Ret1^, and H2B^Htb1^ were adapted from Ling et al., 2024^32^ (n = number of trajectories).

## Notes

### Competing Interest Statement

The authors have declared no competing interest.

### Summary of Updates

Figure scale bars revised; References revised.

