## Supplementary Table 1 for "Live-cell single-molecule dynamics of eukaryotic RNA polymerase machineries"

### Table of contents

**Supplementary Table 1.** Yeast strains for this study.

| **Name** | **Genotype** |
|---|---|
| Parent | *MATa his3-Δ1 leu2-Δ1 trp1-Δ63 ura3-52* |
| *pdr5Δ* | *MATa his3-Δ1 leu2-Δ1 trp1-Δ63 ura3-52 pdr5Δ::loxP* |
| GGEGp | *MATa his3-Δ1 leu2-Δ1 trp1-Δ63 ura3-52 pdr5Δ::loxP GAR1-GFP(S65T)-TRP1 ELO3-GFP(S65T)-natMX6* |
| GGEGp CTD9 | *MATa his3-Δ1 leu2-Δ1 trp1-Δ63 ura3-52 pdr5Δ::loxP GAR1-GFP(S65T)-TRP1 ELO3-GFP(S65T)-natMX6 RPO21-nCTD9* |
| Rpo21-Halo | *GGEGp RPO21-nCTD26-Halo* |
| Rpa190-Halo | *GGEGp RPA190-Halo-kanMX6* |
| Ret1-Halo | *GGEGp RET1-Halo-kanMX6* |
| Rrn9-Halo | *GGEGp RRN9-Halo-kanMX6* |
| Rrn6-Halo | *GGEGp RRN6-Halo-kanMX6* |
| Rrn3-Halo | *GGEGp RRN3-Halo-kanMX6* |
| Pzf1-Halo | *GGEGp PZF1-Halo-kanMX6* |
| Bdp1-Halo | *GGEGp BDP1-Halo-kanMX6* |
| Tfc4-Halo | *GGEGp TFC4-Halo-kanMX6* |
| Tfc3-Halo | *GGEGp TFC3-Halo-kanMX6* |
| Halo-Spt15 | *GGEGp Halo-SPT15* |
| Halo-Toa1 | *GGEGp Halo-TOA1* |
| Sua7-Halo | *GGEGp SUA7-Halo-kanMX6* |
| Taf1-Halo | *GGEGp TAF1-Halo-kanMX6* |
| Rgr1-Halo | *GGEGp RGR1-Halo-kanMX6* |
| Med1-Halo | *GGEGp MED1-Halo-kanMX6* |
| Tfa1-Halo | *GGEGp TFA1-Halo-kanMX6* |
| Tfg1-Halo | *GGEGp TFG1-Halo-kanMX6* |
| Tfb4-Halo | *GGEGp TFB4-Halo-kanMX6* |
| Kin28-Halo | *GGEGp KIN28-Halo-kanMX6* |
| Mot1-Halo | *GGEGp MOT1-Halo-kanMX6* |
| Ncb2-Halo | *GGEGp NCB2-Halo-kanMX6* |
| Cyc8-Halo | *GGEGp CYC8-Halo-kanMX6* |
| Reb1-Halo | *GGEGp REB1-Halo-kanMX6* |
| Spt7-Halo | *GGEGp SPT7-Halo-kanMX6* |
| Spt8-Halo | *GGEGp SPT8-Halo-kanMX6* |
| Halo-Dst1 | *GGEGp Halo-DST1* |
| Hpr1-Halo | *GGEGp HPR1-Halo-kanMX6* |
| Spt16-Halo | *GGEGp SPT16-Halo-kanMX6* |
| Spt4-Halo | *GGEGp SPT4-Halo-kanMX6* |
| Spt6-Halo | *GGEGp SPT6-Halo-kanMX6* |
| Spn1-Halo | *GGEGp SPN1-Halo-kanMX6* |
| Cet1-Halo | *GGEGp CET1-Halo-kanMX6* |
| Abd1-Halo | *GGEGp ABD1-Halo-kanMX6* |
| Sto1-Halo | *GGEGp STO1-Halo-kanMX6* |
| Prp40-Halo | *GGEGp PRP40-Halo-kanMX6* |
| Prp39-Halo | *GGEGp PRP39-Halo-kanMX6* |
| Npl3-Halo | *GGEGp NPL3-Halo-kanMX6* |
| Ctk1-Halo | *GGEGp CTK1-Halo-kanMX6* |
| Sgv1-Halo | *GGEGp SGV1-Halo-kanMX6* |
| Ess1-Halo | *GGEGp ESS1-Halo-kanMX6* |
| Set1-Halo | *GGEGp SET1-Halo-kanMX6* |
| Set2-Halo | *GGEGp SET2-Halo-kanMX6* |
| Set3-Halo | *GGEGp SET3-Halo-kanMX6* |
| Rco1-Halo | *GGEGp RCO1-Halo-kanMX6* |
| Pcf11-Halo | *GGEGp PCF11-Halo-kanMX6* |
| Hrp1-Halo | *GGEGp HRP1-Halo-kanMX6* |
| Cft1-Halo | *GGEGp CFT1-Halo-kanMX6* |
| Ssu72-Halo | *GGEGp SSU72-Halo-kanMX6* |
| Rtt103-Halo | *GGEGp RTT103-Halo-kanMX6* |
| Nrd1-Halo | *GGEGp NRD1-Halo-kanMX6* |
| Halo-Htb1 | *GGEGp Halo-HTB1 PUS1-yomiRFP670nano3-hphMX6* |
| Halo-Htb1, Halo-Htb2 | *GGEGp Halo-HTB1 Halo-HTB2* |
| Halo-Hht1 | *GGEGp Halo-HHT1* |
| Nsi1-Halo | *GGEGp NSI1-Halo-kanMX6* |
| Rpo21-nCTD9-Halo | *GGEGp RPO21-nCTD9-Halo* |

| | |
|---|---|
| Halo-Spt15 CTD9 | *GGEGp CTD9 Halo-SPT15* |
| Halo-Toa1 CTD9 | *GGEGp CTD9 Halo-TOA1* |
| Sua7-Halo CTD9 | *GGEGp CTD9 SUA7-Halo-kanMX6* |
| Taf1-Halo CTD9 | *GGEGp CTD9 TAF1-Halo-kanMX6* |
| Rgr1-Halo CTD9 | *GGEGp CTD9 RGR1-Halo-kanMX6* |
| Med1-Halo CTD9 | *GGEGp CTD9 MED1-Halo-kanMX6* |
| Tfa1-Halo CTD9 | *GGEGp CTD9 TFA1-Halo-kanMX6* |
| Tfg1-Halo CTD9 | *GGEGp CTD9 TFG1-Halo-kanMX6* |
| Tfb4-Halo CTD9 | *GGEGp CTD9 TFB4-Halo-kanMX6* |
| Kin28-Halo CTD9 | *GGEGp CTD9 KIN28-Halo-kanMX6* |
| Mot1-Halo CTD9 | *GGEGp CTD9 MOT1-Halo-kanMX6* |
| Ncb2-Halo CTD9 | *GGEGp CTD9 NCB2-Halo-kanMX6* |
| Cyc8-Halo CTD9 | *GGEGp CTD9 CYC8-Halo-kanMX6* |
| Reb1-Halo CTD9 | *GGEGp CTD9 REB1-Halo-kanMX6* |
| Spt7-Halo CTD9 | *GGEGp CTD9 SPT7-Halo-kanMX6* |
| Spt8-Halo CTD9 | *GGEGp CTD9 SPT8-Halo-kanMX6* |
| Halo-Dst1 CTD9 | *GGEGp CTD9 Halo-DST1* |
| Hpr1-Halo CTD9 | *GGEGp CTD9 HPR1-Halo-kanMX6* |
| Spt16-Halo CTD9 | *GGEGp CTD9 SPT16-Halo-kanMX6* |
| Spt4-Halo CTD9 | *GGEGp CTD9 SPT4-Halo-kanMX6* |
| Spt6-Halo CTD9 | *GGEGp CTD9 SPT6-Halo-kanMX6* |
| Spn1-Halo CTD9 | *GGEGp CTD9 SPN1-Halo-kanMX6* |
| Cet1-Halo CTD9 | *GGEGp CTD9 CET1-Halo-kanMX6* |
| Abd1-Halo CTD9 | *GGEGp CTD9 ABD1-Halo-kanMX6* |
| Sto1-Halo CTD9 | *GGEGp CTD9 STO1-Halo-kanMX6* |
| Prp40-Halo CTD9 | *GGEGp CTD9 PRP40-Halo-kanMX6* |
| Prp39-Halo CTD9 | *GGEGp CTD9 PRP39-Halo-kanMX6* |
| Npl3-Halo CTD9 | *GGEGp CTD9 NPL3-Halo-kanMX6* |
| Ctk1-Halo CTD9 | *GGEGp CTD9 CTK1-Halo-kanMX6* |
| Sgv1-Halo CTD9 | *GGEGp CTD9 SGV1-Halo-kanMX6* |
| Ess1-Halo CTD9 | *GGEGp CTD9 ESS1-Halo-kanMX6* |
| Set1-Halo CTD9 | *GGEGp CTD9 SET1-Halo-kanMX6* |
| Set2-Halo CTD9 | *GGEGp CTD9 SET2-Halo-kanMX6* |
| Set3-Halo CTD9 | *GGEGp CTD9 SET3-Halo-kanMX6* |
| Rco1-Halo CTD9 | *GGEGp CTD9 RCO1-Halo-kanMX6* |
| Pcf11-Halo CTD9 | *GGEGp CTD9 PCF11-Halo-kanMX6* |
| Hrp1-Halo CTD9 | *GGEGp CTD9 HRP1-Halo-kanMX6* |
| Cft1-Halo CTD9 | *GGEGp CTD9 CFT1-Halo-kanMX6* |
| Ssu72-Halo CTD9 | *GGEGp CTD9 SSU72-Halo-kanMX6* |
| Rtt103-Halo CTD9 | *GGEGp CTD9 RTT103-Halo-kanMX6* |
| Nrd1-Halo CTD9 | *GGEGp CTD9 NRD1-Halo-kanMX6* |
| Halo-Htb1 CTD9 | *GGEGp CTD9 Halo-HTB1 PUS1-yomiRFP670nano3-hphMX6* |

**Supplementary Table 1.** Yeast strains for this study.
